# The effect of PEGylation on surface tethering of liposomes via DNA nanotechnology

**DOI:** 10.1101/2025.07.07.663613

**Authors:** James Gaston, Sreelakshmi Meepat, Md Sirajul Islam, Jasleen Kaur Daljit Singh, Michael J Booth, Shelley FJ Wickham, Matthew AB Baker

## Abstract

Polyethylene glycol (PEG) is widely used in liposome formulation due to its blocking properties and ability to prolong circulation *in vivo*, to create biomimetic liposomes and drug delivery devices. Similarly, membrane-embedded DNA nanotechnology is increasingly used to modulate cellular behaviour and communication. However, there is a gap in knowledge in how PEG-lipid formulations can be optimised for both liposome properties and control of selective DNA hybridisation. To address this, we systematically investigated the effect of liposome PEG content on DNA mediated tethering of liposomes to glass surfaces. We formulated liposomes of two different lipid compositions (DOPE/DOPC or DPhPC), with varying amounts of PEGylated lipid (0-50%). We measured the effect of increased PEG content on liposome size and polydispersity through dynamic light scattering (DLS). Small amounts of PEG (0-20%) introduced repulsive forces that reduced size, while large amounts of PEG (30-50%) increased polydispersity. PEG-liposomes were then decorated with cholesterol-DNA strands and labelled with either intercalating lipid dyes or fluorescently labelled lipids. Binding to surfaces via complementary DNA strands was quantified using total internal reflection fluorescence (TIRF) microscopy. We found that PEGylation of DNA-liposomes could either block or enhance surface binding, depending on the amount of PEG. DNA-liposomes with reduced surface binding included DPhPC/DiD with 10% or 20% PEG-lipid. In contrast, DNA-liposome surface binding increased for DOPE/DOPC/DiD with increasing PEG%. This study highlights that while PEG can act to stabilise liposome formulations, its ability to block specific DNA binding interactions on membranes is variable and dependent on membrane composition.

**Table of contents figure:** 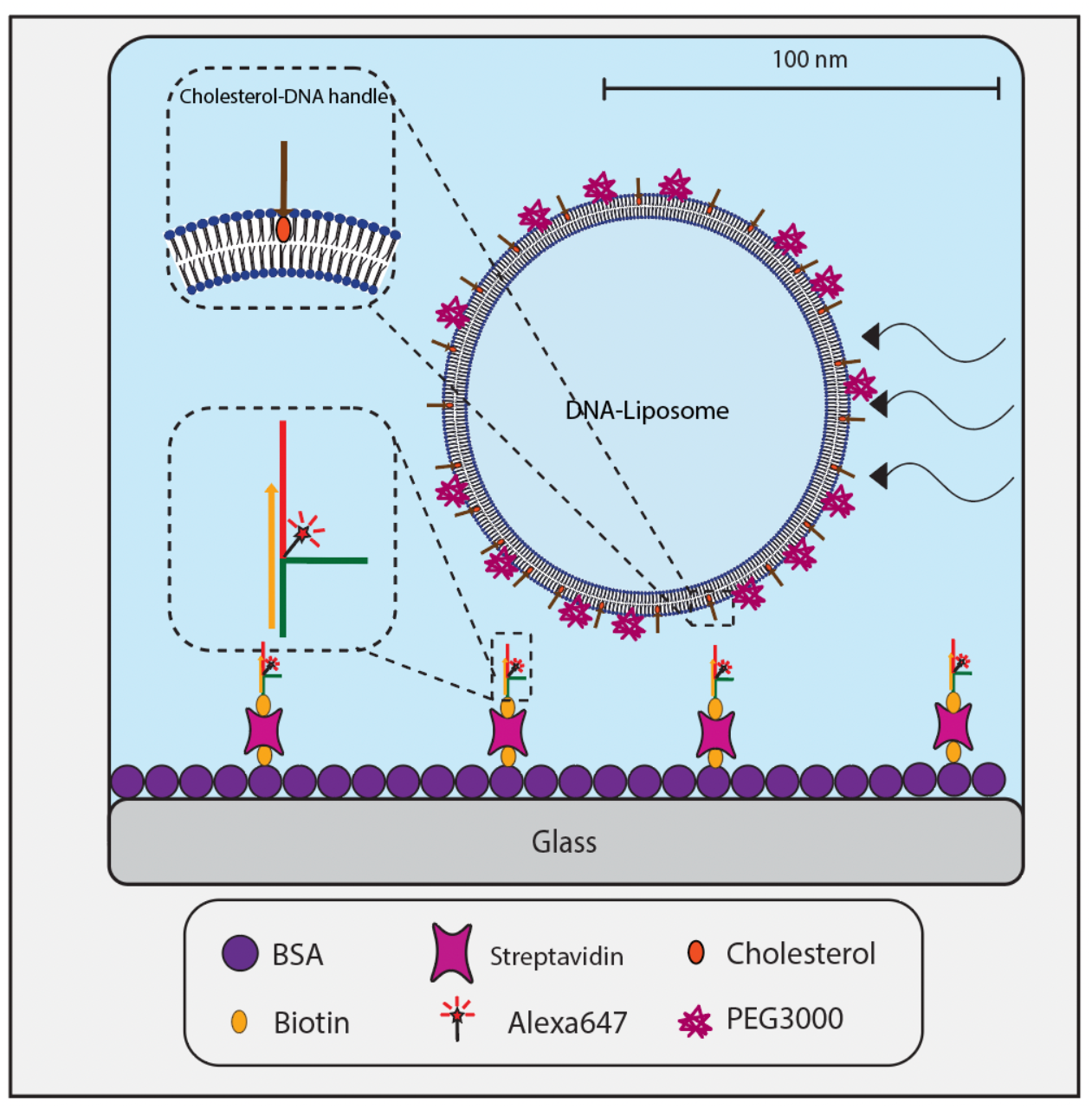

## Introduction

PEG is a hydrophilic polymer primarily known for its surface blocking capabilities (D’souza and Shegokar, 2016). It is often incorporated with other membrane components to encapsulate therapeutic payloads. Notably, PEG is used as a component of lipid nanoparticles (LNPs) to evade immune system responses and prevent cellular uptake to improve circulation times *in vivo* (Hatakeyama *et al*., 2011). PEG has also been used to mediate membrane interactions, including creating more stable membranes in droplet systems (Booth *et al*., 2016), and as a blocker of non-specific surface fouling for molecules like DNA, proteins, and lipids (Liu *et al*., 2013; Downs *et al*., 2020). Addition of PEG has improved the design of functionalised liposomes that can incorporate targeting groups, such as aptamers, to direct and control specific membrane interactions. The capacity of PEG to block non-specific interactions has been studied (Karimata, Nakano and Sugimoto, 2007), and there is growing interest in the role of PEG in aiding precise control of membrane interactions to functionalise membranes effectively (Wu *et al*., 2014).

DNA can be used as a programmable and addressable building block to self-assemble 2D and 3D biomimetic nanostructures through complementary base pairing (Seeman, 1982). Liposomes can be decorated with DNA nanostructures via cholesterol moieties (Ohmann et al., 2019). Such technology integrated with membranes has broadened the *in vivo* and *ex vivo* applications of liposomes. Programmable, complementary base pairing of DNA can direct binding interactions between membranes, allowing for tethering of membranes together and on surfaces at controllable distances (Löffler, Ries and Vogel, 2020).

Previous studies have shown that PEG can assist in directing membranes through blocking non-specific aggregation and binding (Shi *et al*., 2021; Xu *et al*., 2023). It remains to be shown whether PEG can be useful in blocking the specific interactions of DNA functionalised membranes that are energetically driven to interact by DNA hybridisation. Quantifying this mechanism of blocking is a key component in expanding future design capabilities of DNA functionalised liposomes as vehicles for delivering encapsulated payloads. Here we investigate the effect of increased concentration of PEG-lipid on the formulation of DNA-coated liposomes and quantify the disruption of specific surface binding interactions mediated by DNA hybridisation.

## Materials and methods

### DNA complex synthesis

Sequences for all DNA strands and complexes (Supplementary Table 1) were generated using NUPACK design software (Zadeh *et al*., 2011) to test for the formation of unwanted secondary structures. DNA strands were purchased pre-modified with appropriate moieties (Integrated DNA Technologies, Inc., USA). The assembly reaction of the DNA complex was prepared by mixing the ssDNA at a ratio of 1:1 with a final concentration of 5 µM in 1xTAE buffer (Tris-base 5 mM, EDTA 1 mM). The mixture of ssDNA was incubated overnight at room temperature under constant motion.

### Gel formation and PAGE analysis

DNA complexes assembled under different conditions and formations were observed using non-denaturing polyacrylamide gel electrophoresis (PAGE). Polyacrylamide gels were formed using polyacrylamide that was at a 30% stock concentration with a 37.5:1 mono:bis acrylamide ratio (Bio-Rad Laboratories, USA). For observation of annealed complexes, 5.33 mL of the polyacrylamide was used in a 10 mL total solution to form a 16% polyacrylamide gel. The remainder of the solution was filled with a combination of 200 μL of a 50x stock of Tris-Acetate-EDTA (TAE) buffer, 5 μL of N,N,N′,N′ -Tetramethyl ethylenediamine (TEMED), 80 μL of 10% Ammonium persulfate (APS) in water, and MilliQ water to have a final solution volume of 10 mL. Once the gel solution was mixed, 8 mL of solution was added to SureCast™ gel casting plates (Invitrogen Inc., USA) to be cast using the SureCast™ gel casting module (Invitrogen Inc., USA). 15 well gel combs were used, and gels were left to polymerise overnight at 4°C before use.

Preformed DNA complexes were premixed in tubes with MilliQ water to ensure there was 150 ng of total DNA. 1 μL of 5x loading dye was added to this DNA mix to make a final volume of 5 μL per tube. DNA and loading dye mixtures were added to each lane of the cast gel and run in 1x TAE buffer on the Invitrogen Mini Gel Tank Electrophoresis system (Invitrogen Inc., USA) connected to a Lively 300V Power Supply (Major Science Co., Ltd, China). Gels were run at 150 V over 90 min. 5 μL of 10000x concentrated SYBR™ Gold Nucleic Acid Gel Stain (Invitrogen Inc., USA) was added to 50 mL MilliQ with the gel submerged under constant shaking to post-stain for 4 min. Gels were imaged using Invitrogen™ iBright™ FL1500 Imaging System (Thermo Fisher Scientific Inc., USA).

### Liposome preparation

Liposomes were produced by extrusion using a Mini-Extruder (Avanti Polar Lipids Inc., USA). Lipid mixtures of DOPE and DOPC lipids (Avanti Polar Lipids Inc., USA) at a 1:1 ratio, or DPhPC lipid (Avanti Polar Lipids Inc., USA) were dissolved in chloroform at a 10 mg/mL concentration. 18:1 PEG3000-PE PEGylated lipid (Avanti Polar Lipids Inc., USA) was added as a percentage of this initial 10 mg/mL lipid mixture where appropriate. When labelling for fluorescence imaging, lipid mixtures where combined with 0.1% molar mass of the lipophilic carbocyanine dye DiD (Thermo Fisher Scientific Inc., USA), or PE-Rhodamine lipids (Avanti Polar Lipids Inc., USA).

100 μL of lipid solution was added to a round-bottom glass tube and dried at room temperature under nitrogen gas flow while constantly rotating the tube, creating a thin, uniform coating of lipids over the bottom 20 mm of the tube. Extrusion buffer (100 mM NaCl, 10 mM MgCl2, 5 mM Tris-HCl, pH 7.5) was added to the tube post-drying to suspend the lipids at a concentration of 1 mg/mL. The tube was then sealed with Parafilm (Bemis Company, Inc., USA), vortex mixed at room temperature for 3 minutes, and sonicated at 30°C in a water bath sonicator for 10 minutes to create a turbid aqueous suspension of lipids.

The lipid suspension was transferred to a 500 μL glass syringe (Hamilton Company, UK) and passed back and forth through a 100 nm polycarbonate filter (Whatman plc, USA) 41 times to produce a suspension of extruded homogenous liposomes. Polycarbonate filters, along with filter supports (Avanti Polar Lipids Inc., USA) were replaced between extrusions of the two fluorescent species to prevent cross-contamination of fluorescence. Liposomes were diluted 100-fold in extrusion buffer to form a liposome suspension of a final concentration of 0.01 mg/mL in preparation for fluorescence imaging. Where appropriate, cholesterolated ssDNA handle strands with varying concentrations (Supplementary Table 1) were incubated for 1 hour on a rotating carousel with liposomes to incorporate into the membrane bilayer while limiting unwanted aggregation.

### Dynamic Light Scattering (DLS) characterisation of liposome size with PEG3000 formulations

Liposome size measurements were performed using a Zetasizer Nano ZS system (Malvern Instruments, Worcestershire, UK) equipped with a 633 nm He–Ne laser and operating at an angle of 173°. After extrusion, liposomes (with membrane tethered PEG and DNA where appropriate) were diluted to imaging concentration (0.01 mg/mL) in extrusion buffer to a total volume of 1 mL and loaded into a disposable cuvette for size measurement. All measurements were performed in triplicate. Measurements were carried out using the following Standard Operating Procedure (SOP) settings: Material (Phospholipids); Material Refractive index (1.45); Dispersant Viscosity (0.8659 cP); Dispersant Refractive Index (1.332). Data was acquired and analysed using Zetasizer Software (V8.01, Malvern). These measurements gave final outputs in the form of size distribution graphs, which included distribution curves representing how size varies across a liposome population, quantification of what liposome size represented the peak of these distributions, and the standard deviation for these distribution curves.

### Imaging and Microscopy

Tunnel slides for imaging of DNA-liposome interaction were constructed using a 50 mm cover slip (#1 thickness) (Menzel Glaser GmbH) that was fixed to a glass slide (Suzhou Upline Medical Products Co., China (PRC)) using two parallel strips of double-sided tape (Nichiban Co., Japan) approximately 150 μm thick placed 4 mm apart, forming a thin channel of approximately 20 μL volume (a ‘tunnel’ slide). A thin layer of CoverGrip Coverslip Sealant (Biotium Inc., USA) was applied over remaining exposed tape to prevent the contamination of solutions by adhesive residue and left to dry for 5 minutes. Solution exchange was executed by adding solution to one end of the channel with a pipette while simultaneously drawing solution from the opposite end with an absorbent paper wipe (Kimberly-Clark Professional, USA).

To block the glass coverslip surface, 30 μL of a 9:1 mixture of bovine serum albumin (BSA) and biotinylated bovine serum albumin (BSA-biotin) in MilliQ water at 0.5 mg/mL total concentration was added to the slide and incubated for 10 minutes. This allowed the blocking of non-specific binding of DNA and fluorescent liposomes to the glass, while also enabling biotinylated components to be tethered to the surface via biotin-streptavidin-biotin conjugation. Excess BSA and BSA-biotin in solution was then removed by flushing 40 μL of the appropriate buffer through the slide (extrusion buffer for liposome imaging or 1X TAE buffer for surface DNA imaging). To conjugate BSA-biotin to biotin moieties on either DNA or liposomes, 30 μL of streptavidin at 0.5 mg/mL in MilliQ water was subsequently flowed in and incubated for 10 minutes. To prevent streptavidin-mediated aggregation, unbound streptavidin was washed out by flushing 40 μL of the extrusion buffer or 1X TAE buffer through the slide. Biotinylated DNA was then added at 30 μL volume to the slide and incubated for 10 minutes in order to tether to the slide surface via biotin-streptavidin-biotin conjugation. To remove unbound DNA, slides were washed using 40 μL of extrusion buffer. Finally, 30 μL of fluorescent DNA-liposomes in extrusion buffer was added to the slide and incubated for 10 min to allow hybridisation between surface-bound DNA constructs and the DNA-liposomes in solution. After incubation, slides were washed using 40 μL of extrusion buffer to remove unbound DNA-liposomes.

Slides were imaged using Total Internal Reflection Fluorescence (TIRF) imaging on a Zeiss Elyra PALM Superresolution Microscope with a 63x/1.4 Oil Iris M27 oil immersion objective (Carl Zeiss AG, Germany) and Andor iXon 897 EMCCD camera (Oxford Instruments plc, United Kingdom). For each slide, five images were taken of arbitrarily selected points around the centre of the tunnel slide, selecting areas with uniformly dispersed liposomes or DNA. For imaging fluorescently labelled liposomes, a 642 nm laser was used at 10% power (0.0376 µW/µm^2^ sample exposure) for DiD labelled liposomes, and a 561 nm laser was used at 10% power (0.2204 µW/µm^2^ sample exposure) for Rhodamine labelled liposomes. For Alexa647 labelled DNA, a 642 nm laser at 30% power (0.1128 µW/µm^2^ sample exposure) was used for imaging. All 642 nm laser images were taken using a 655 nm long pass emission dichroic filter while 546 nm laser images were taken using a 535/40 nm band pass together with a 750 nm long pass dichroic filter. Camera integration time of 33 ms and line averaging of two was used for all images, and laser power was selected in order to match intensities of fluorophore emission output.

### Image Analysis

Fluorescence images were analysed using custom image analysis workflow (Supplementary Figure 3) to identify fluorescence regions of interest (ROIs) and extract information to characterise surface binding. Briefly, source images were split into two fluorescence channels, and the channel of interest was converted into a pixel value array. The array was thresholded based on a predetermined value to separate signal from background and form a binary array of pixels above the threshold value. The threshold was set based upon assessment of image background for our specific imaging set-up, with minimum fluorescence values being 2 standard deviations (calculated across all pixels) above the maximum value of background fluorescence in a blank microscopy image. To quantify ROIs, contours were applied to outline the edges of areas of the image above the threshold. For fluorescently labelled DNA samples, the average number of DNA fluorescence ROIs identified above the threshold was denoted as *N*_spots_. For liposomes, once contours were outlined, they were further separated based on an area threshold into two categories, liposome spot contours and aggregate contours. Aggregates were determined to be any contour larger than 50 pixels in total area (Supplementary Figure 3). The number of liposome spots (*N*_spots_) and aggregates (*N*_agg_) were normalised against the average number in the 0% peg condition using the following equations:

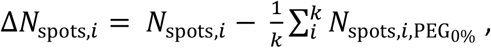

where for image *i*, of a set of images *k*, the change in number of liposome spot contours (Δ*N*_spots,*i*_) is calculated as the number of liposome contours in a specific image after subtracting the mean number of liposome spot contours over all images with no PEG3000 (PEG_0%_).

Likewise for Δ*N*_agg_:

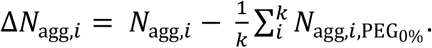

For aggregate contours, the change in total pixel area (Δ*A*_agg_) for all aggregates in image *i* of a set *k* was determined similarly as:

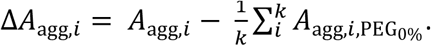

## Results

We designed a modular DNA complex to act as a tether for liposome surface binding (Figure 1A and B). Formation of the complex was confirmed through PAGE analysis, with the complex forming a band with mobility equivalent to a 300 bp duplex. This was slower than the mobility expected from the 63 bp length (Supplementary Figure 1). The DNA complex was subsequently modified with a biotin moiety for surface immobilisation via biotin-streptavidin surface chemistry. An Alexa647 fluorophore modification was also added for visualisation through TIRF microscopy (Figure 1C). The complex was confirmed to be surface bound via TIRF microscopy and the surface density of single fluorophore DNA regions of interest (ROIs) was found to increase in a concentration dependent manner with the addition of DNA complex (Figure 1D).

**Figure 1.**
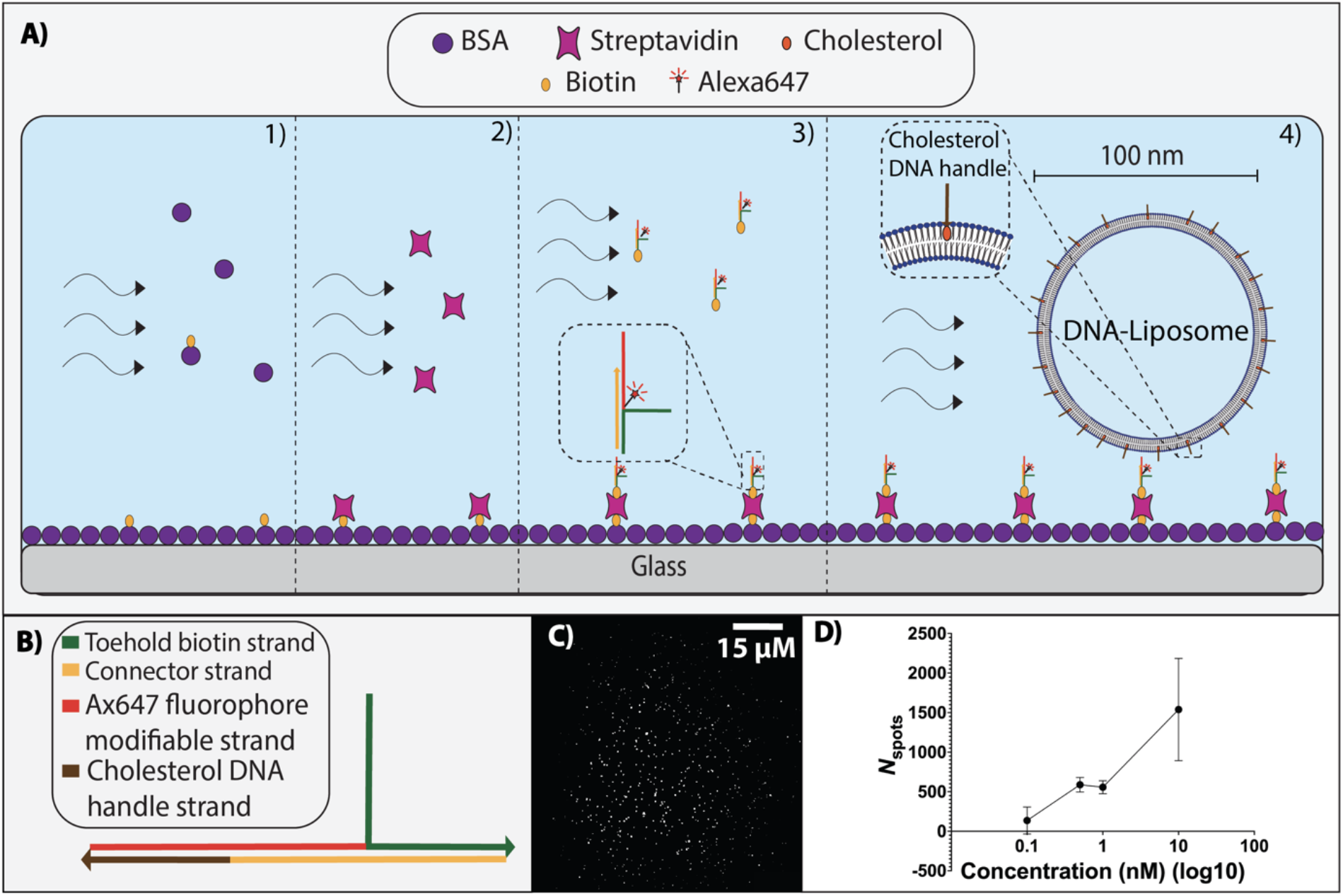
Characterisation of DNA complex shown to bind to BSA blocked glass surfaces through biotin-streptavidin binding. **A)** Stepwise schematic for surface blocking and immobilisation of Alexa647 labelled DNA complex. 1) BSA-biotin is flowed in, then binds streptavidin (2). Biotinylated-DNA complex is added (3), and lastly liposomes coated in complementary ssDNA are flowed in, which then surface tether via DNA hybridisation of complementary ssDNA on the liposome and the surface complex. **B)** Schematic representation of DNA complex with labelled DNA components. Each line colour corresponds to a different DNA strand denoted in the figure legend. Arrows indicate 5’ to 3’ direction of DNA sequence. **C)** Representative TIRF microscopy image of 1 nM biotin-DNA-Alexa647 complex surface-immobilised through biotin-streptavidin binding, scale bar 15um. **D)** The number of identified DNA binding events (*N*_spots_) for the biotinylated Alexa647 DNA complex increases as surface incubation concentration increases.

Surface tethering of liposomes was characterised by incubating fluorescent DNA-liposomes, pre-labelled with cholesterol DNA handle strands, with surface-immobilised biotin-DNA complex. Hybridisation of the cholesterol DNA handle strand (chol-DNA handles) to the surface DNA complex was shown to be sequence specific. The number of surface ROIs (*N*_spots,_), indicative of the number of liposomes bound to the surface, was 806 ± 645 ROIs when 50 nM of biotinylated DNA complex was used to coat the surface. At the same concentration of a strand designed to have a scrambled, non-complementary sequence (Supplementary Table 1) a lower *N*_spots_ of 63 ± 10 ROIs was observed (Figure 2B). The fully complementary sequence therefore had an approximately 12-fold increase in surface binding compared with a scrambled sequence. Surface binding was found to increase with the concentration of biotin-DNA complex immobilised on the surface (Figure 2B). Increasing the concentration of liposome anchored cholesterol-DNA handles also increased surface binding when the surface concentration of biotin-DNA complex was kept constant (1 nM) (Figure 2C). Greatest surface binding (661 ± 222 ROIs) occurred when 10 nM of cholesterol-DNA was pre-incubated to decorate liposomes (0.01 mg/mL lipid concentration) prior to surface tethering (Figure 2C).

**Figure 2.**
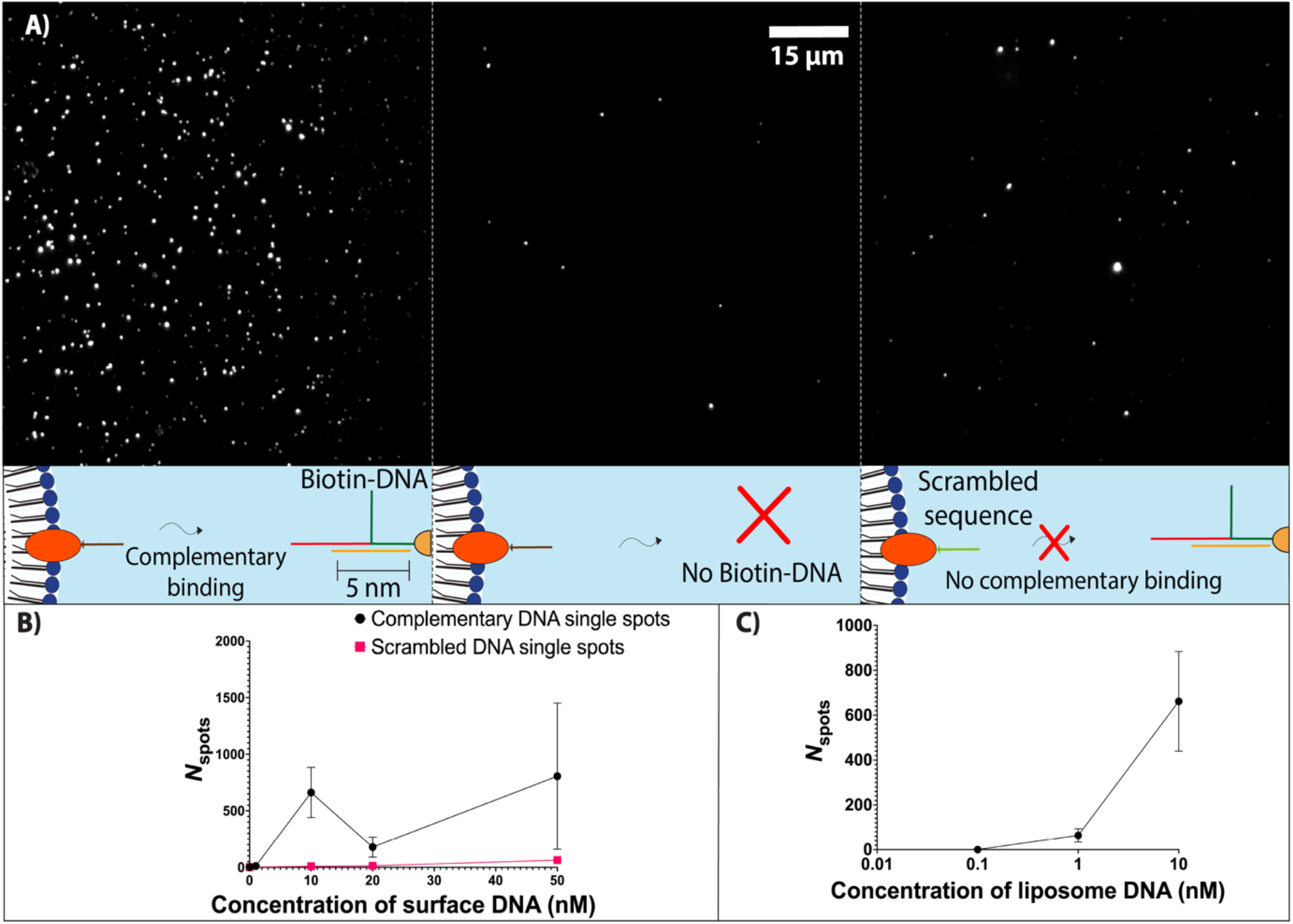
Sequence specific hybridisation of cholesterolated DNA (chol-DNA) and surface-immobilised biotin-DNA complex increases surface tethering. **A)** Representative images and schematic representations of DiD labelled fluorescent liposomes decorated with 10 nM complementary cholesterol-DNA handles (left), no DNA handles (middle), and 10 nM scrambled (non-complementary) cholesterol-DNA handles (right). **B)** Number of fluorescence spots above background (*N*_spots_) is used as a measure of the number of DNA-liposomes bound to the surface, and increases as concentration of complementary surface DNA (biotinylated DNA complex) is increased. Concentration of liposome anchored cholesterol-DNA handles (complementary or scrambled sequences) is held constant at 10 nM. **C)** *N*_spots_ increases as concentration of liposome DNA (complementary cholesterol-DNA handles decorating liposomes) is increased. Concentration of surface-immobilsed biotinylated DNA is held constant at 1 nM.

PEG3000 conjugated phospholipid (PEG3000-PE) was incorporated into the liposome formulation at increasing amounts. DLS was used to validate the size of liposomes for both DPhPC and DOPE/DOPC lipid formulations, reporting the distribution of sizes, size peaks, and polydispersity index (PdI) of each liposome population (Figure 3). As the percentage of PEG-lipid increased, DPhPC liposomes remained at a consistent size (between 117 and 122 d.nm) but polydispersity increased from 0.136 to 0.246 PdI (Figure 3A and B). For DOPE/DOPC liposomes, the size and polydispersity initially decreased between 0% and 20% PEG3000, with a 8 d.nm decrease in size and 0.089 PdI decrease in polydispersity. This was followed by increase in polydispersity up to 0.23 PdI with 40% PEG3000 (Figure 3C and D). DLS characterisation indicated that the 3 PEG-lipid formulations that had the lowest liposome polydispersity were 0%, 10%, and 20% PEG3000-PE for both lipid compositions.

**Figure 3.**
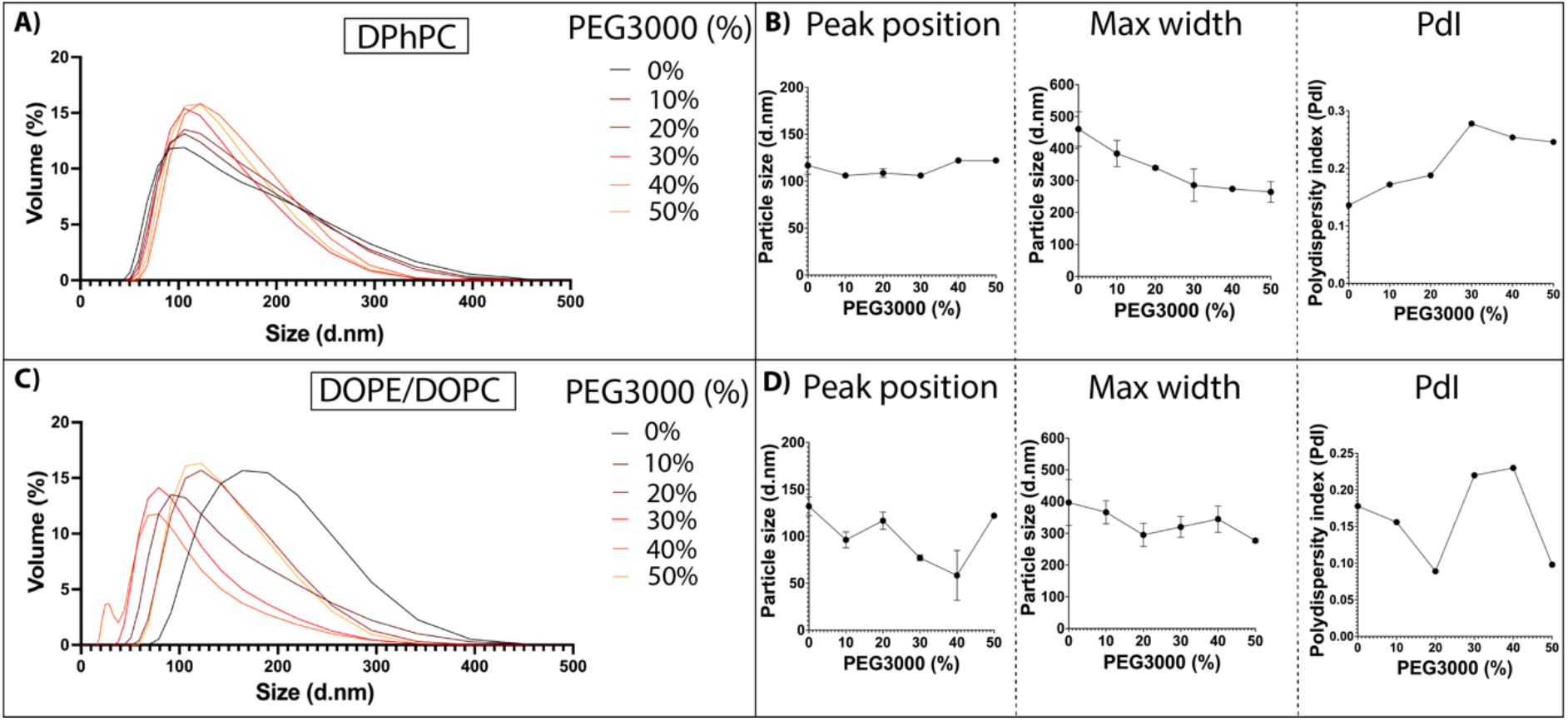
DLS characterisation of liposome formulations with varying amounts of PEG3000 indicates changes in size and polydispersity. **A)** DLS line plots showing size distributions for DPhPC liposomes with 0-50% PEG3000 added to the initial lipid mix. **B)** Position of the DLS curve peak (left), along with the max width (width from smallest to largest size measured) of the DLS curves (middle) and polydispersity index (right) for DPhPC liposomes plotted against the percentage of PEG3000 in liposome formulation. **C)** DLS line plots showing size distributions for DOPE/DOPC liposomes with 0-50% PEG3000 added to the initial lipid mix. **D)** Position of the DLS curve peak (left), along with the max width of the DLS curves (middle) and polydispersity index (right) for DOPE/DOPC liposomes plotted against the percentage of PEG3000 in liposome formulation.

The three lipid formulations of 0%, 10%, and 20% PEG3000 were subsequently used to assess surface binding interactions mediated by DNA hybridisation interactions. Membrane bound cholesterol-DNA handles and surface immobilised complementary biotin-DNA complex were kept at fixed concentrations of 10 nM and 1 nM, respectively. DiD was initially used as an intercalating fluorescent dye to label both DPhPC and DOPE/DOPC liposomes (Figure 4A and B). For DPhPC liposomes significant differences in Δ*N*_spots_ were observed with changing PEG formulation (p<0.05). Δ*N*_spots_ progressively decreased from 0 ± 8 to -51 ± 2 ROIs (p<0.001) between 0% and 20% PEG3000 formulation (Figure 4C). There was no significant difference in Δ*N*_agg_ between 0% and 10% PEG3000 formulation, but there was a significant decrease in Δ*N*_agg_ (p<0.001) between 10% (10 ± 12 ROIs, p) and 20% (−23 ± 1 ROIs) PEG3000 formulation (Figure 4C). There was no significant variation for Δ*A*_agg_ between 0% and 10% PEG3000 formulation, but there was a significant decrease in Δ*A*_agg_ (p<0.05) between 10% (917 ± 2659 contour area) and 20% (−3284 ± 38 pixels) PEG3000 formulation (Figure 4D). There was also an overall decrease in fluorescent pixel brightness from 9882 ± 814 AU to 6687 ± 768 AU between 0% to 20% PEG3000 formulation (Figure 4E).

**Figure 4.**
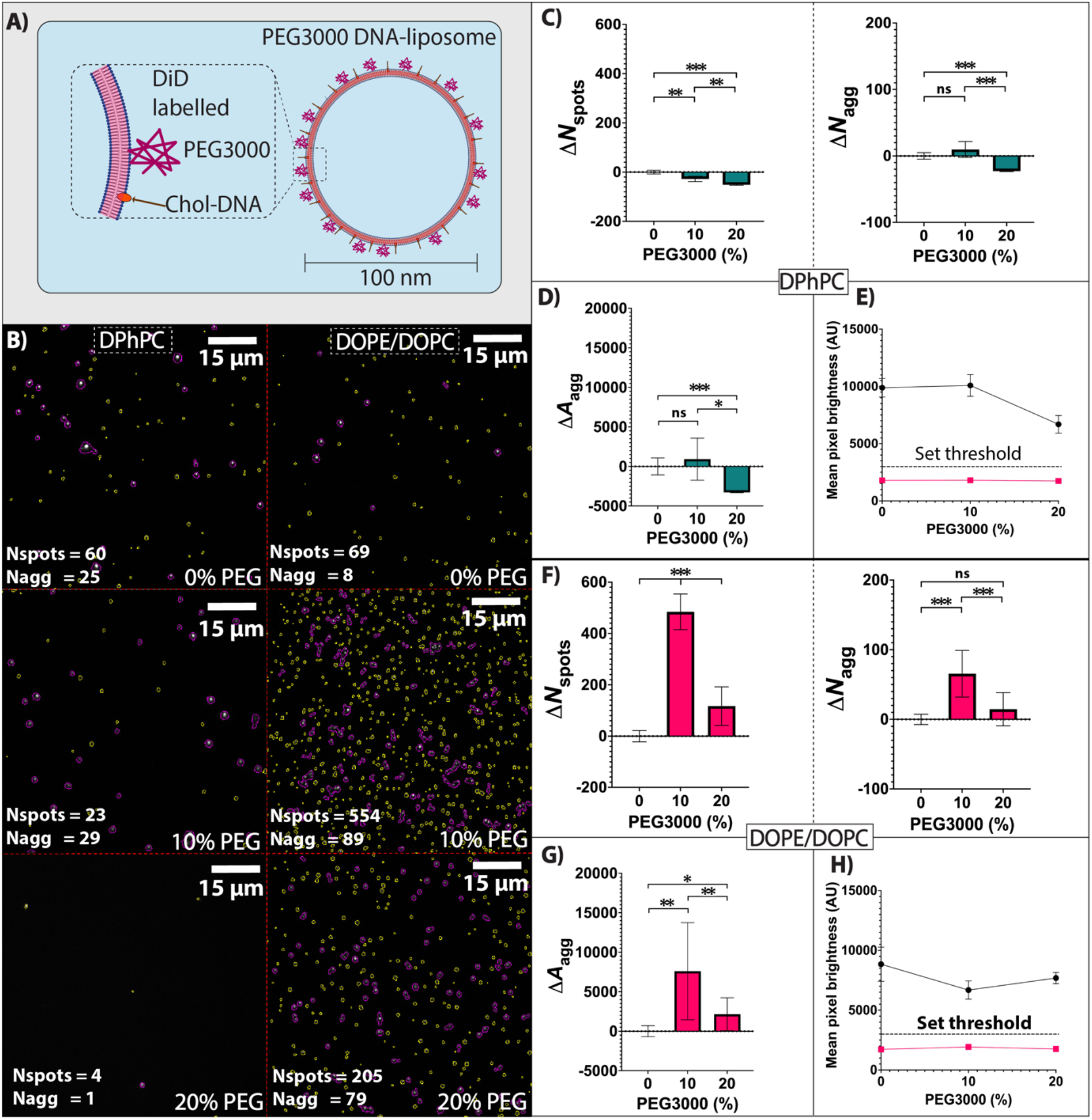
Differing PEG3000 levels in formulation of DiD labelled DNA-liposomes interferes with surface tethering. **A)** Schematic showing the fluorescence labelling for PEGylated DNA-liposomes with DiD. **B)** Representative microscopy images showing the different levels of surface binding between DiD labelled DPhPC (left) and DOPE/DOPC (right) liposomes with 0% (top), 10% (middle) and 20% (bottom) PEG3000 being incorporated in liposome formulation. Liposome spot ROIs are highlighted in yellow, and aggregate ROIs are highlighted in magenta. The number of liposome spot ROIs (*N*_spots_) and aggregate ROIs (*N*_agg_) are listed on the bottom left of each representative image. **C)** Change in surface binding for DiD labelled DPhPC liposomes for liposome spot ROIs (Δ*N*_spots_, left) and aggregate ROIs (Δ*N*_agg_, right) with increasing PEG3000 percentage. **D)** Change in total pixel area of all aggregate ROIs (Δ*A*_agg_) and **(E)** comparison of mean pixel brightness above (black) and below (magenta) the set fluorescence threshold (dotted line, 3000 AU). This threshold was set to separate background from liposome signal for DPhPC. **F)** Change in surface binding through for DiD labelled DOPE/DOPC liposomes measured for liposome spot ROIs (Δ*N*_spots_, left) and aggregate ROIs (Δ*N*_agg_, right) with increasing PEG3000 percentage. **G)** Change in total pixel area of all aggregate ROIs (Δ*A*_agg_) and **(H)** comparison of mean pixel brightness above (black) and below (magenta) the set fluorescence threshold (dotted line, 3000 AU). This threshold was set to separate background from liposome signal for DOPE/DOPC. For all p-values, ns = p≥0.05, * = p<0.05, ** = p<0.01, and *** = p<0.001.

For DOPE/DOPC liposomes there was significant difference (p<0.05) for both Δ*N*_spots_ and Δ*N*_agg_. For Δ*N*_spots_ there was an increase from 0 ± 22 to 484 ± 69 ROIs (p<0.001) between 0% and 10% PEG3000 formulation, followed by a decrease to 117 ± 75 ROIs (p<0.001) for 20% PEG3000 formulation (Figure 4F). For Δ*N*_agg_ there was a similar increase from 0 ± 7 to 66 ± 33 (p<0.001) between 0% and 10% PEG3000 formulation, followed by a decrease to 15 ± 24 ROIs (p<0.001) for 20% PEG3000 formulation (Figure 4F). There was a significant increase in Δ*A*_agg_ (p<0.01) between 0% (0 ± 697 pixels) and 10% (7593 ± 6150 pixels) PEG3000 formulation, followed by a significant decrease (p<0.01) for 20% PEG3000 formulation (2152 ± 2079 pixels) (Figure 4G). There was also a decrease in pixel brightness from 8845 ± 1431 AU to 6681 ± 767 AU from 0% to 10% PEG3000 formulation, followed by an increase to 7685 ± 469 AU for 20% PEG3000 formulation (Figure 4H).

To assess how the method of labelling affected surface binding, the type of fluorophore labelling was changed from an intercalating fluorescent dye (DiD) to a fluorescently labelled lipid (PE-Rhodamine). For DPhPC liposomes, the variation in Δ*N*_spots_ was not significant but there was a significant increase (p<0.001) from 0 ± 24 to 67 ± 11 ROIs in Δ*N*_agg_ for 10% PEG3000 formulation, followed by a decrease (p<0.001) to -33 ± 7 ROIs in Δ*N*_agg_ for 20% PEG3000 formulation (Figure 5C). There was no significant variation for Δ*A*_agg_ between 0% and 10% PEG3000 formulation, but there was a significant decrease in Δ*A*_agg_ (p<0.001) between 10% (5362 ± 8146 pixels) and 20% (−21746 ± 1643 pixels) PEG3000 formulation (Figure 5D). There was also an overall decrease in fluorescent pixel brightness from 11044 ± 2748 AU to 8225 ± 1457 AU between 0% to 20% PEG3000 formulation (Figure 5E).

**Figure 5.**
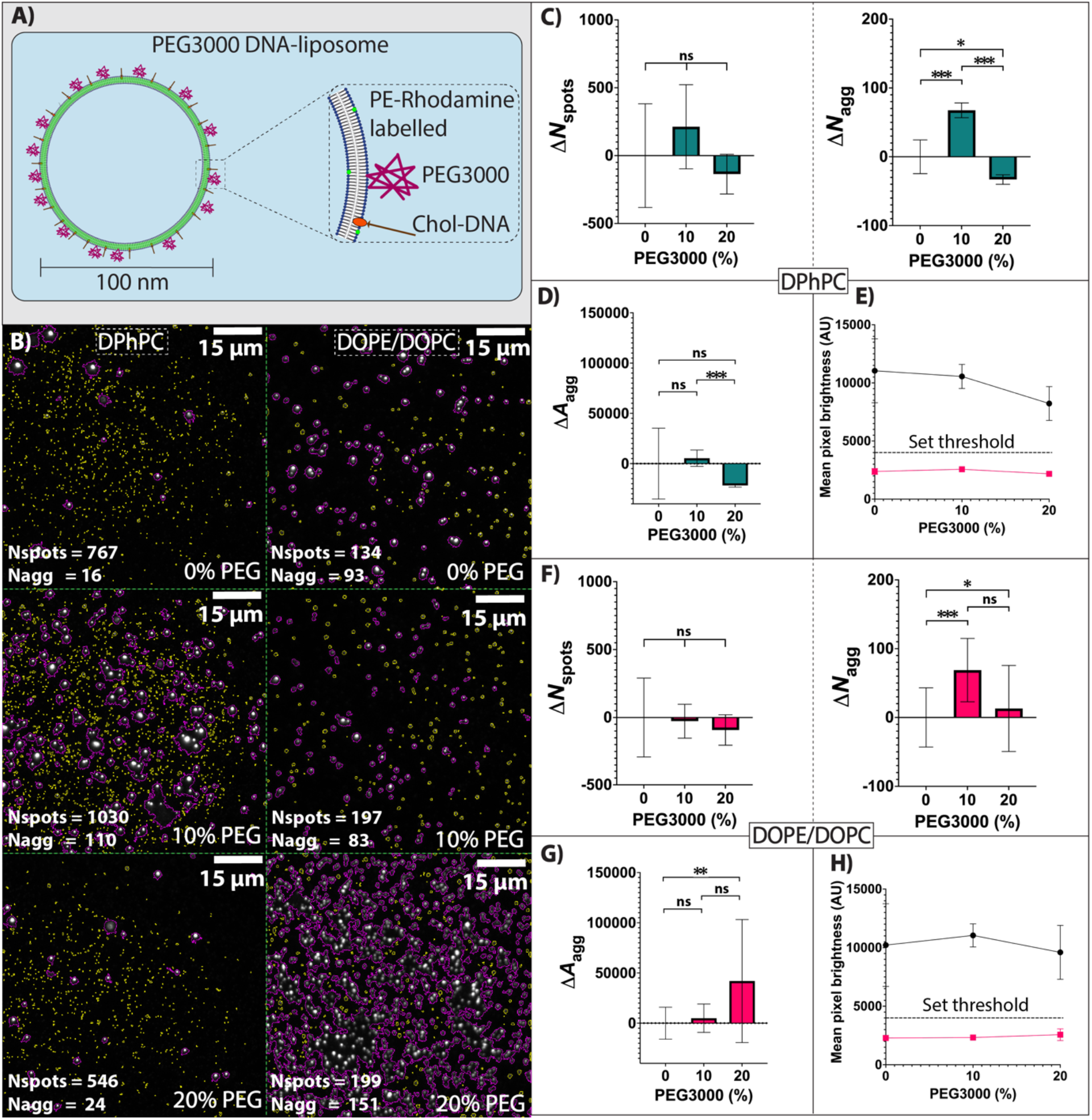
Differing PEG3000 levels in formulation of PE-Rhodamine labelled DNA-liposomes interferes with surface tethering. **A)** Schematic showing the fluorescence labelling for PEGylated DNA-liposomes with PE-Rhodamine. **B)** Representative microscopy images showing the different levels of surface binding between PE-Rhodamine labelled DPhPC (left) and DOPE/DOPC (right) liposomes with 0% (top), 10% (middle) and 20% (bottom) PEG3000 being incorporated in liposome formulation. Liposome spot ROIs are highlighted in yellow, and aggregate ROIs are highlighted in magenta. The number of liposome spot ROIs (*N*_spots_) and aggregate ROIs (*N*_agg_) are listed on the bottom left of each representative image. **C)** Change in surface binding for PE-Rhodamine labelled DPhPC liposomes for liposome spot ROIs (Δ*N*_spots_, left) and aggregate ROIs (Δ*N*_agg_, right) with increasing PEG3000 percentage. **D)** Change in total pixel area of all aggregate ROIs (Δ*A*_agg_) and **(E)** comparison of mean pixel brightness above (black) and below (magenta) the set fluorescence threshold (dotted line, 4000 AU). This threshold was set to separate background from liposome signal for DPhPC. **F)** Change in surface binding for PE-Rhodamine labelled DOPE/DOPC liposomes for liposome spot ROIs (Δ*N*_spots_, left) and aggregate ROIs (Δ*N*_agg_, right) with increasing PEG3000 percentage. **G)** Change in total pixel area of all aggregate ROIs (Δ*A*_agg_) and **(H)** comparison of mean pixel brightness above (black) and below (magenta) the set fluorescence threshold (dotted line, 4000 AU). This threshold was set to separate background from liposome signal for DOPE/DOPC. For all p-values, ns = p≥0.05, * = p<0.05, ** = p<0.01, and *** = p<0.001.

For DOPE/DOPC liposomes labelled with PE-rhodamine, the variation in Δ*N*_spots_ was not significant, but there was an increase (p<0.001) from 0 ± 43 to 69 ± 46 ROIs in Δ*N*_agg_ for 10% PEG3000 formulation (Figure 5F). There was no significant variation in Δ*N*_agg_ between 10% and 20% PEG3000 formulation (Figure 5F). There was no significant variation for Δ*A*_agg_ between 0% and 10%, and 10% to 20% PEG3000 formulation. There was a significant increase in Δ*A*_agg_ (p<0.01) between 0% (0 ± 15895 pixels) and 20% (41958 ± 61256 pixels) PEG3000 formulation (Figure 5G). There was an increase in pixel brightness from 10219 ± 3535 AU to 11048 ± 987 AU from 0% to 10% PEG3000 formulation, followed by a decrease to 9601 ± 2301 AU for 20% PEG3000 formulation (Figure 5H).

## Discussion

### DNA complex characterisation

PAGE characterisation of our DNA complex indicated a size of approximately 300 bp compared with an expected size of 63 bp. This larger size was likely due to the branched ssDNA toehold section of the complex causing a slower migration than a purely double stranded DNA duplex of the same base pair length (Stellwagen and Stellwagen, 2020). Conjugation of liposomes to glass coverslips aligned with expectations from previous studies that used DNA binding interactions to immobilise DNA-liposomes onto surfaces (Malle *et al*., 2022). This was validated through reduced surface binding when scrambled non-complementary cholesterol-DNA handles were used.

### Characterisation of PEG incorporation and its role in membrane formation

Since PEG was added as part of the initial lipid mix before liposome formulation, the amount of membrane PEG may be heterogeneous between individual liposomes. DLS characterisation and PdI values indicated that when liposome formulation included over 30% PEG3000-PE, particle size and population heterogeneity increased suggesting an uneven distribution of PEG incorporation across the population of liposomes.

Comparing 0% and 20% PEG3000 formulations, PdI values were low and size distribution peaks decreased by 25.8 d.nm for DOPE/DOPC liposomes, and 17.5 d.nm for DPhPC liposomes. This reduction in size has been previously described as being due to the slightly negative charge of PEG-lipid promoting lateral repulsion forces, resulting in the curving of the bilayer and causing a slight reduction in liposome diameter (Sriwongsitanont and Ueno, 2004; Tsermentseli *et al*., 2018; Jaradat, Meziane and Lamprou, 2024).

This reduction in size was not as noticeable with DPhPC bilayers, suggesting that the effect of membrane PEG3000 incorporation in DPhPC bilayers differs from DOPE/DOPC bilayers. Membrane properties like fluidity, thickness, and density of lipid packing could account for such variation. DPhPC membranes are more tightly packed and may maintain shape and uniformity when formulated with PEG compared with DOPC/DOPE membranes which are more fluid (Reddy, Warshaviak and Chachisvilis, 2012; Kara *et al*., 2017).

### Role of membrane composition in PEG mediated binding

We used TIRF microscopy to further quantify individual liposome surface interactions. We tested two types of lipid compositions (DPhPC, and a mix of DOPE/DOPC), two different types of fluorescent labelling (intercalating DiD and fluorescent lipid PE-Rhodamine) and a range of membrane formulations with increasing percentage of a linear PEG type (PEG3000). Our hypothesis was that an increasing percentage of PEG would reduce surface binding by blocking DNA hybridisation interactions. However, we observed that the relationship was not linear and increased PEG did not always result in reduced surface binding.

We separated the data into 2 populations: surface bound liposomes (*N*_spots_) and surface bound aggregates (*N*_agg_) to more accurately determine surface binding effects on individual liposomes. PE-Rhodamine labelled DOPE/DOPC and DPhPC liposomes had no significant difference in Δ*N*_spots_ across PEG lipid formulations, but an increase in Δ*N*_agg_ at 10% PEG lipid formulation. PEG was also shown to reduce surface binding for DiD labelled DPhPC liposomes with increasing PEG formulations, while DiD labelled DOPE/DOPC liposomes had both increased Δ*N*_spots_ and Δ*N*_agg_ with increasing PEG content. This aggregation was not detected in solution measurements, suggesting that DLS is not as adept at detecting small aggregation events. Aggregation on coverslip surfaces for different membrane formulations is likely due to enriched surface binding of aggregates with multiple DNA handles. Aggregation was most notable in conditions like PE-Rhodamine DOPE/DOPC liposomes with 20% PEG lipid formulation, and the aggregation in these conditions could affect measurement accuracy as aggregate areas can mask nearby individual liposome spots. Careful assessment of optimal liposome concentration may mitigate this effect.

One factor that could affect liposome surface interactions is the PEG type – whether branched or linear. Linear PEG types have been shown to have non-linear increases in viscosity with increasing concentration (Kirinčič and Klofutar, 1999). Other PEG types, such as branched, Y-shaped, and multi-arm PEGs (D’souza and Shegokar, 2016; Xu *et al*., 2023; Bento *et al*., 2024). display different characteristics of size and stability, with branched PEGs known to be more stable than linear PEGs of similar sizes (Fee, 2007). PEGs of increasing molecular weight have been shown to increase steric hinderance due to increased chain length and overall size (Godinho *et al*., 2014). Blocking could thus potentially be improved by varying the type of PEG molecule and increasing molecular weight, which could reduce the variability observed with linear PEG3000 in this study.

Different labels and fluorophores also had a pronounced effect on binding. DiD is an intercalating carbocyanine dye that can incorporate differently within tightly packed DPhPC membranes compared to the more fluid DOPE/DOPC membranes (Axelrod, 1979; Hsieh *et al*., 1997). DiD also has a head group that can be attracted to glass surfaces. If surface blocking via BSA was not complete then there is the potential for increased surface binding with higher levels of DiD (Jeyachandran *et al*., 2009). Fluorescently labelled lipids like PE-Rhodamine mix with other lipids and do not intercalate like the carbocyanine dyes, however the addition of this lipid type can alter overall membrane characteristics (Kleusch *et al*., 2012).

### Future directions

This study highlights the varied impact of PEG content on liposome-surface interactions. Our results indicate that PE-Rhodamine labelled DNA-liposomes with no PEG had the highest proportion of specifically bound individual liposomes when compared with all conditions. DiD labelled DNA-liposomes formulated with DPhPC lipids had the highest reduction in surface interactions as the percentage of PEG3000-PE was increased. This was shown within a narrow set of membrane formulations, however, and innovations in methods for automation of liposome production could allow for further investigation of a large number of membrane compositions in a short time span (Challita *et al*., 2018; Dupin *et al*., 2022; Mason, Wickham and Baker, 2024). Use of such automation and high throughput screening could facilitate optimal searching of a large set of possible conditions to identify the formulation with the desired blocking effects. Future work could also incorporate additional DNA functionality, such as strand displacement activity through ssDNA toehold elements, to direct to where DNA-liposomes should tether or where they are blocked (Löffler *et al*., 2017; Löffler, Ries and Vogel, 2020; Malle *et al*., 2022).

