## Supplementary Figures 1-7, Supplementary Tables 1-4 for "The effect of PEGylation on surface tethering of liposomes via DNA nanotechnology"

### Supplementary information

| DNA strand name | DNA strand sequence | Potential DNA strand modifications |
|---|---|---|
| **(1)** Toehold strand | CCTACTGACTTTATCCACCGATTCTAGGGTTAAAAGGGGACG | 3’ Biotin-TEG |
| **(2)** Connector strand | CGTCCCCTTTTAACCCTAGAAGGGATAAGTTGATTGCAGAGC | None |
| **(3)** Alexa647 fluorophore modifiable strand | TCTCGACACAAATCTTCCTGCGCTCTGCAATCAACTTATCCC | 3’ Alexa Fluor 647 (NHS Ester) |
| **(4)** Chol-DNA handle strand | GCAGGAAGATTTGTGTCGAGA | 3’ Cholesterol-TEG |
| **(5)** Scrambled chol-DNA handle strand | CTGGTGAGACGCTAGAC | 3’ Cholesterol-TEG |

**Supplementary table 1. DNA strand names, sequences, and modifications.** DNA strand names, sequences, and potential strand modifications that can be added to that ssDNA component to add functionality are listed.

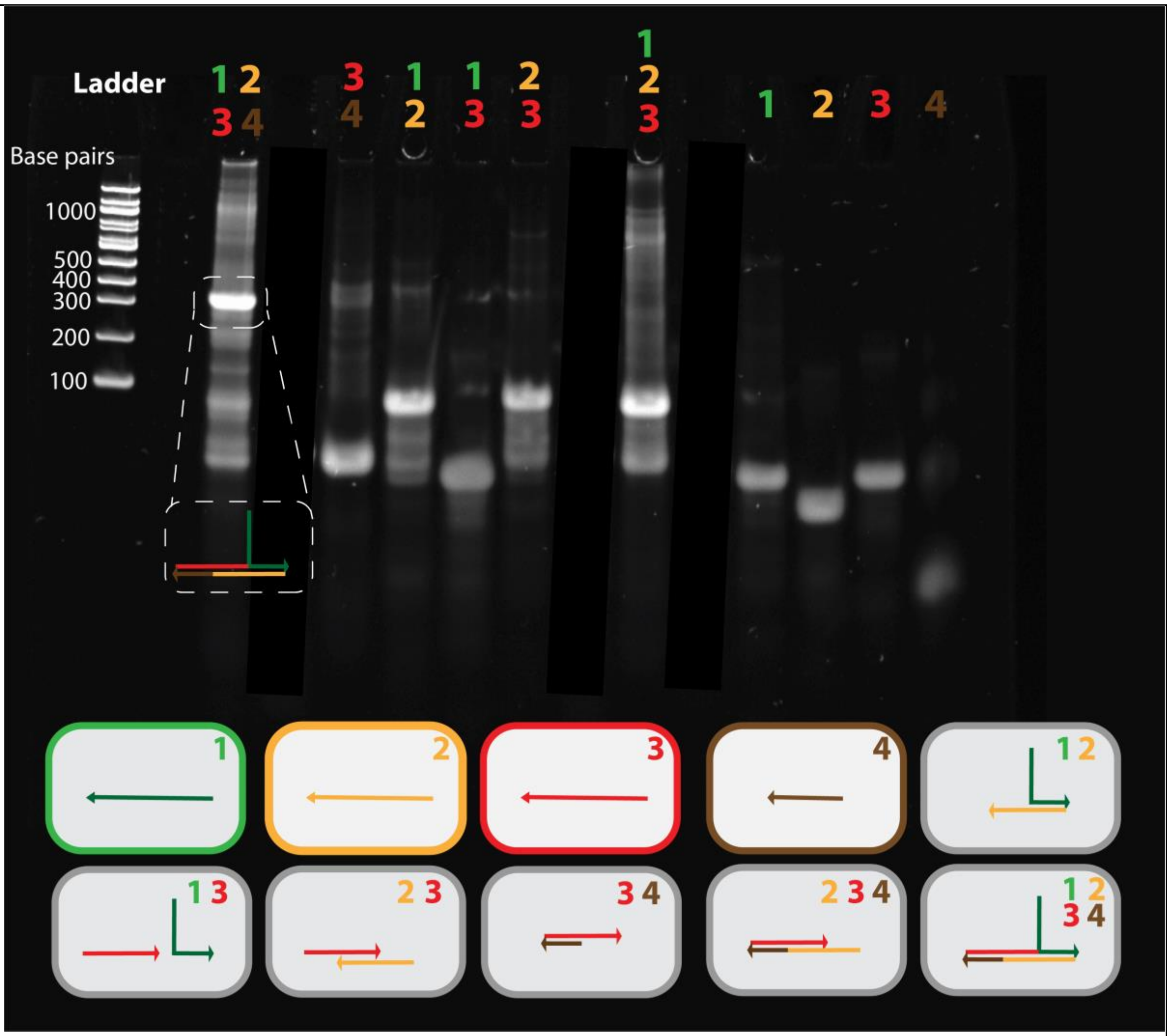


**Supplementary Figure 1. 16% PAGE confirming complex formation.** Individual strands and combinations present within each lane are indicated by numbers (top) corresponding to the legend (bottom). Gel visualisation of the different combinations of ssDNA leading up to the hybridisation of the full complex shows size increases that correspond to larger DNA constructs..

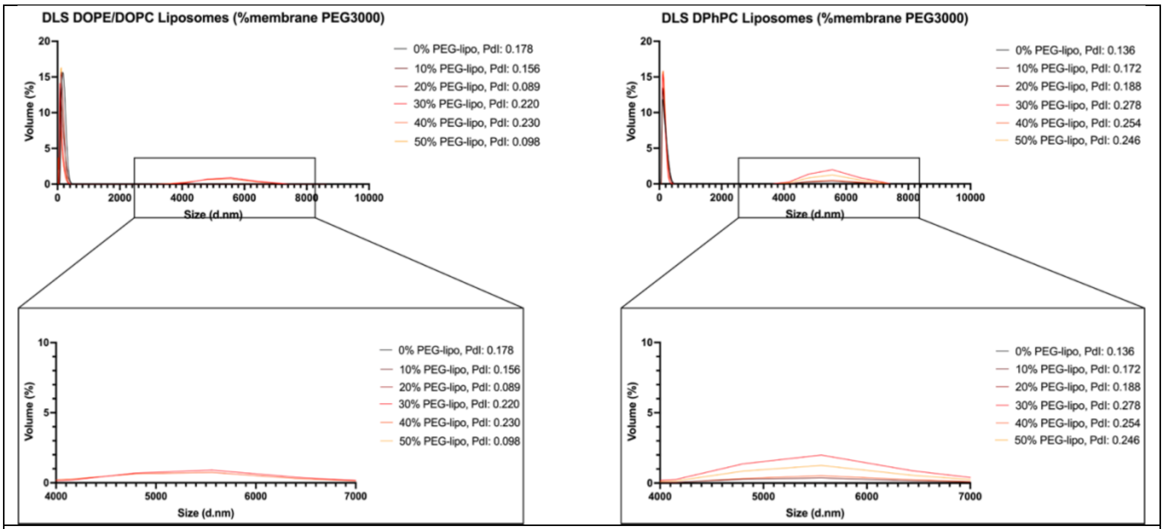


**Supplementary Figure 2. DLS measurements of liposome sizes with varying amounts of PEG present in initial lipid formulations.** Full DLS line plots showing size distributions for both DPhPC (right) and DOPE/DOPC (left) liposomes with 0-50% PEG3000 added to the initial lipid mix formulation. Inset graphs were displayed with different x-axes to show the secondary artefact peaks for each lipid composition (bottom). Polydispersity Index (PdI) values are also included for each condition in the figure legends.

Code available here

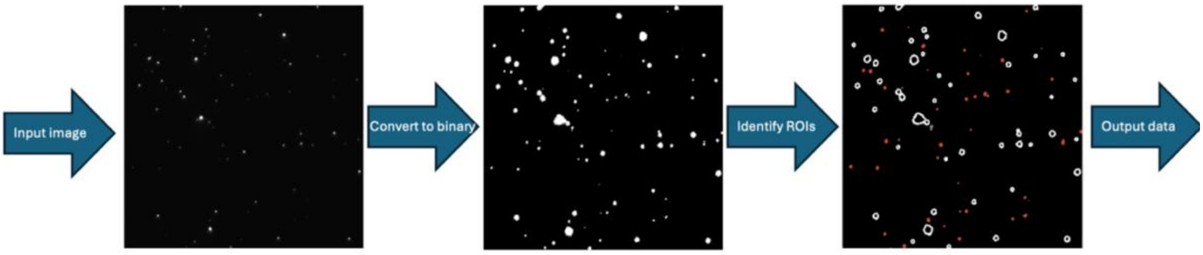


**Supplementary figure 3. Python code and graphical representation of spot detection analysis method for microscopy images.** Analysis code loop was written in Python and run on Python version 3.8. Graphical representation of the image analysis pipeline (bottom) shows how images are processed from raw inputs (left) into binary images based on a minimum intensity threshold that identifies pixels associated with fluorescence ROIs to minimise background interference (middle). This is followed by ROIs being segregated into liposome spots below 50 pixels (right- red outlines) and aggregates above 50 pixels (right- white outlines) based on a chosen area threshold in order to only quantify blocking of individual liposomes as opposed to large aggregates. This fluorescence ROI data is then outputted into an excel file.

| DiD labelled liposomes | | | | |
|---|---|---|---|---|
| **Lipid type** | **PEG3000-PE (%)** | **Image no.** | $N_{spots}$ | $N_{agg}$ |
| **DPhPC** | **0** | #1 | 43 | 16 |
| | | #2 | 60 | 21 |
| | | #3 | 60 | 25 |
| | | #4 | 62 | 27 |
| | | #5 | 56 | 28 |
| | | **Total** | | |
| | | **5** | **281** | **117** |
| | **10** | #1 | 46 | 25 |
| | | #2 | 31 | 35 |
| | | #3 | 20 | 24 |
| | | #4 | 23 | 29 |
| | | #5 | 22 | 53 |
| | | **Total** | | |
| | | **5** | **142** | **166** |
| | **20** | #1 | 3 | 1 |
| | | #2 | 8 | 0 |
| | | #3 | 4 | 1 |
| | | #4 | 4 | 0 |
| | | **Total** | | |
| | | **4** | **19** | **2** |
| **DOPE/DOPC** | **0** | #1 | 69 | 8 |
| | | #2 | 62 | 8 |
| | | #3 | 20 | 6 |
| | | #4 | 16 | 1 |
| | | #5 | 37 | 6 |
| | | #6 | 47 | 26 |
| | | #7 | 59 | 13 |
| | | #8 | 73 | 17 |
| | | #9 | 79 | 11 |
| | | #10 | 65 | 19 |
| | | **Total** | | |
| | | **10** | **527** | **115** |
| | **10** | #1 | 413 | 37 |
| | | #2 | 519 | 75 |
| | | #3 | 598 | 48 |
| | | #4 | 476 | 134 |
| | | #5 | 501 | 106 |
| | | #6 | 554 | 89 |
| | | #7 | 599 | 115 |
| | | #8 | 632 | 64 |
| | | #9 | 487 | 36 |
| | | #10 | 592 | 67 |
| | | **Total** | | |
| | | **10** | **5371** | **771** |
| | **20** | #1 | 205 | 79 |
| | | #2 | 162 | 11 |

| | | | | |
|---|---|---|---|---|
| | | #3 | 163 | 13 |
| | | #4 | 149 | 53 |
| | | #5 | 306 | 34 |
| | | #6 | 204 | 20 |
| | | #7 | 68 | 5 |
| | | #8 | 85 | 12 |
| | | #9 | 254 | 29 |
| | | #10 | 102 | 5 |
| | | | **Total** | |
| | | **10** | **1698** | **261** |

**Supplementary table 2. Number of contour ROIs identified across all DiD labelled liposome conditions.** For DiD labelled liposomes the number of contours was listed for liposome spot ROIs ($N_{spots}$) and aggregate ROIs ($N_{agg}$) for each image, along with the sums of total number of images and total number of ROIs for each PEG3000 liposome formulation.

**PE-Rhodamine labelled liposomes**

| Lipid type | PEG3000-PE (%) | Image no. | $N_{spots}$ | $N_{agg}$ |
|---|---|---|---|---|
| **DPhPC** | **0** | #1 | 1392 | 56 |
| | | #2 | 1121 | 56 |
| | | #3 | 1035 | 30 |
| | | #4 | 484 | 25 |
| | | #5 | 767 | 16 |
| | | #6 | 368 | 73 |
| | | #7 | 585 | 89 |
| | | #8 | 321 | 62 |
| | | #9 | 294 | 44 |
| | | #10 | 443 | 82 |
| | | | **Total** | |
| | | **10** | **6810** | **533** |
| | **10** | #1 | 1248 | 112 |
| | | #2 | 567 | 134 |
| | | #3 | 567 | 130 |
| | | #4 | 1030 | 110 |
| | | #5 | 1057 | 118 |
| | | | **Total** | |
| | | **5** | **4469** | **604** |
| | **20** | #1 | 546 | 24 |
| | | #2 | 353 | 23 |
| | | #3 | 747 | 8 |
| | | #4 | 603 | 24 |
| | | #5 | 475 | 22 |
| | | | **Total** | |
| | | **5** | **2724** | **101** |
| | **0** | #1 | 133 | 102 |
| | | #2 | 994 | 31 |
| | | #3 | 134 | 93 |

| | | | | |
|---|---|---|---|---|
| DOPE/DOPC | | #4 | 293 | 94 |
| | | #5 | 170 | 6 |
| | | #6 | 500 | 9 |
| | | #7 | 350 | 95 |
| | | #8 | 212 | 83 |
| | | #9 | 257 | 1 |
| | | #10 | 1093 | 48 |
| | | #11 | 104 | 0 |
| | | #12 | 175 | 1 |
| | | #13 | 66 | 0 |
| | | #14 | 166 | 0 |
| | | #15 | 42 | 0 |
| | | #16 | 551 | 80 |
| | | #17 | 475 | 84 |
| | | #18 | 304 | 85 |
| | | #19 | 654 | 87 |
| | | #20 | 261 | 97 |
| | | **Total** | | |
| | | **20** | **6934** | **996** |
| | **10** | #1 | 197 | 83 |
| | | #2 | 276 | 83 |
| | | #3 | 455 | 56 |
| | | #4 | 254 | 86 |
| | | #5 | 600 | 81 |
| | | #6 | 335 | 190 |
| | | #7 | 186 | 153 |
| | | #8 | 285 | 141 |
| | | #9 | 272 | 141 |
| | | #10 | 335 | 172 |
| | | **Total** | | |
| | | **10** | **3194** | **1186** |
| | **20** | #1 | 213 | 76 |
| | | #2 | 343 | 146 |
| | | #3 | 199 | 151 |
| | | #4 | 232 | 91 |
| | | #5 | 318 | 127 |
| | | #6 | 373 | 8 |
| | | #7 | 454 | 4 |
| | | #8 | 163 | 5 |
| | | #9 | 103 | 13 |
| | | #10 | 150 | 7 |
| | | **Total** | | |
| | | **10** | **2548** | **628** |

**Supplementary table 3. Number of contour ROIs identified across all PE-Rhodamine labelled liposome conditions.** For PE-Rhodamine labelled liposomes the number of contours was listed for liposome spot ROIs ($N_{spots}$) and aggregate ROIs ($N_{agg}$) for each image, along with the sums of total number of images and total number of ROIs for each PEG3000 liposome formulation.

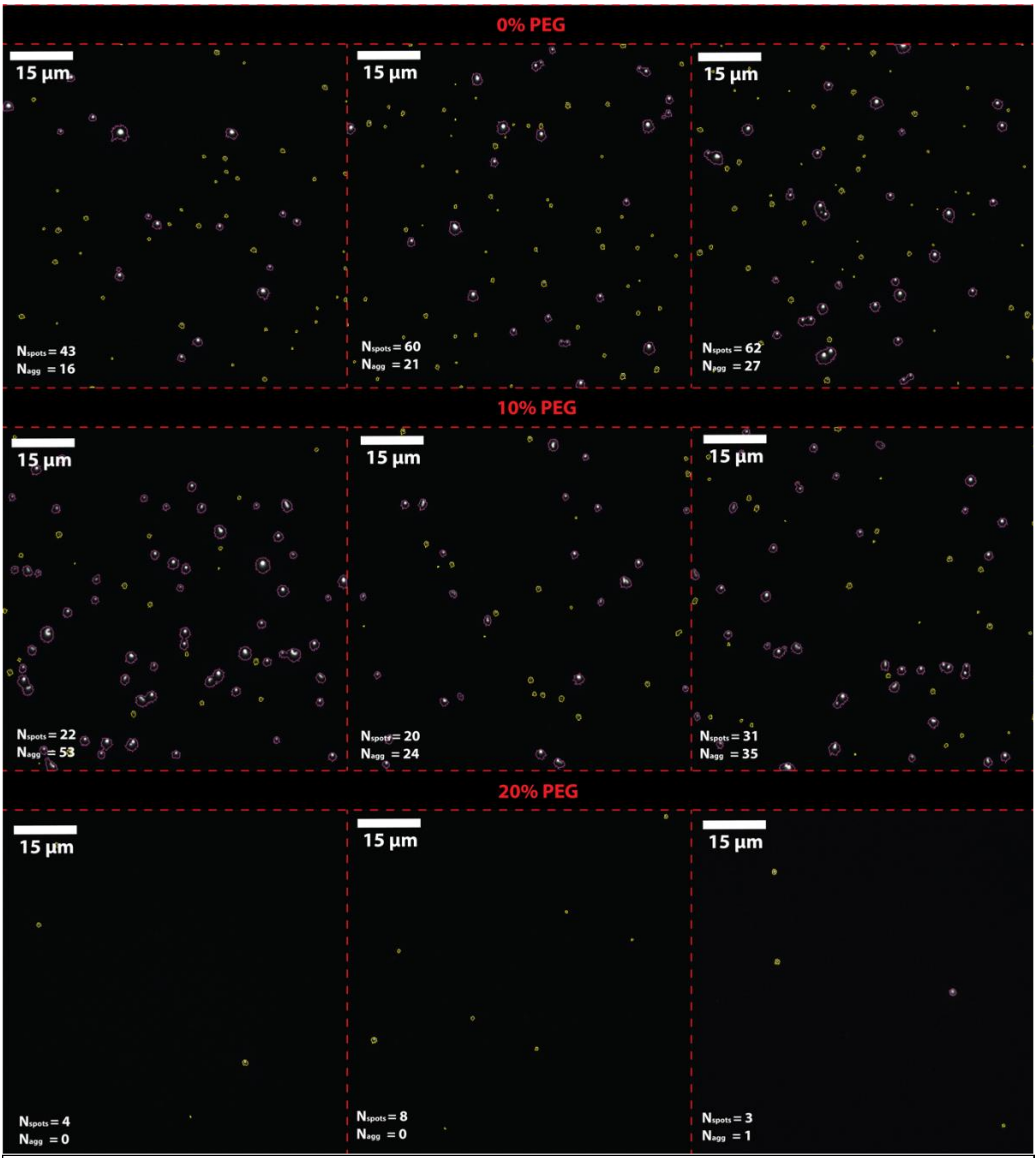


**Supplementary figure 4. Representative TIRF microscopy images of DiD labelled DPhPC liposomes.** Three representative images were chosen for each PEG3000 formulation (0% top, 10% middle, 20% bottom). Liposome spot ROIs were highlighted in yellow, and aggregate ROIs were highlighted in pink. The number of liposome spot ROIs ($N_{spots}$) and aggregate ROIs ($N_{agg}$) are listed on the bottom left of each representative image.

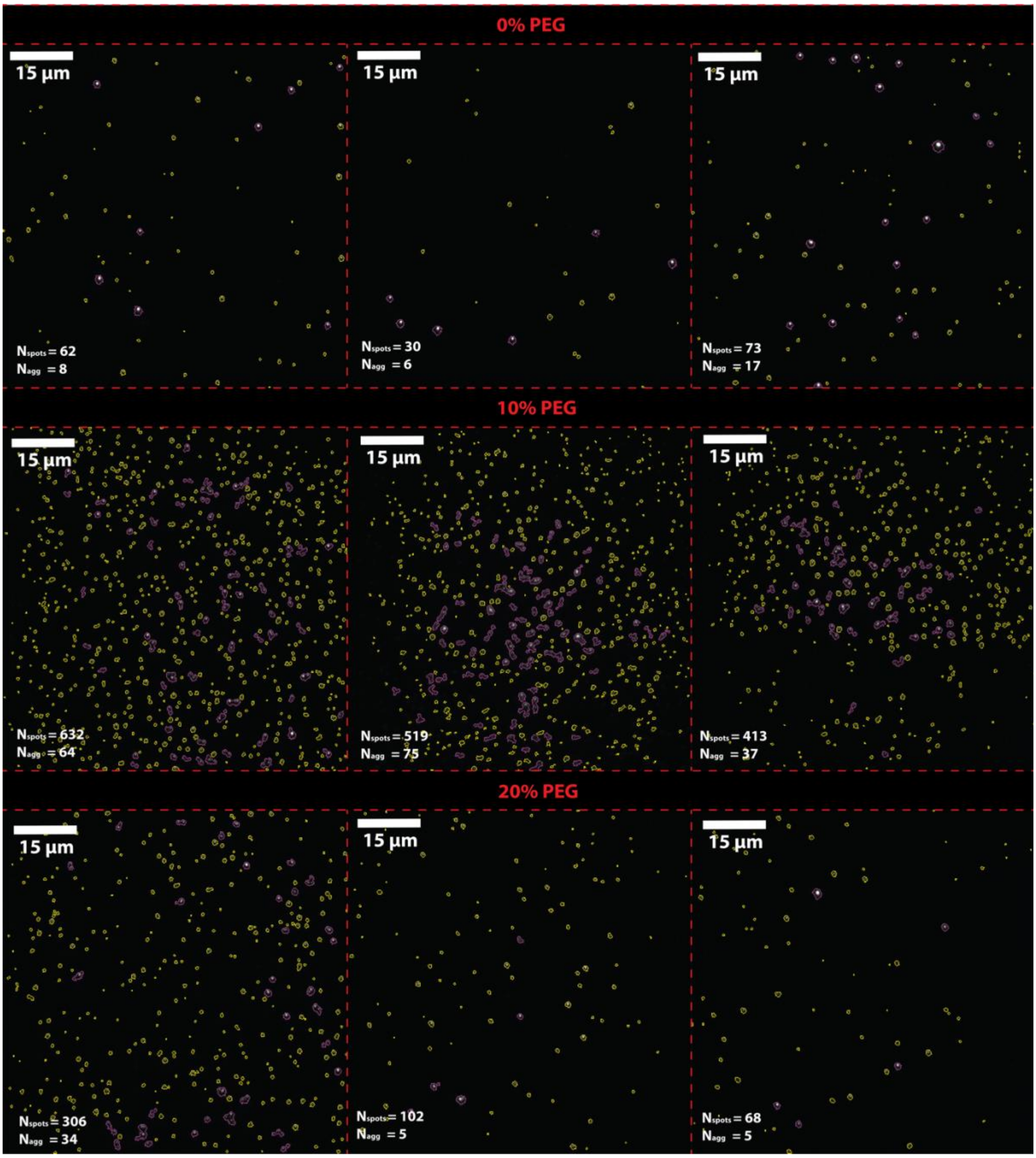


**Supplementary figure 5. Representative TIRF microscopy images of DiD labelled DOPE/DOPC liposomes.** Three representative images were chosen for each PEG3000 formulation (0% top, 10% middle, 20% bottom). Liposome spot ROIs were highlighted in yellow, and aggregate ROIs were highlighted in pink. The number of liposome spot ROIs ($N_{spots}$) and aggregate ROIs ($N_{agg}$) are listed on the bottom left of each representative image.

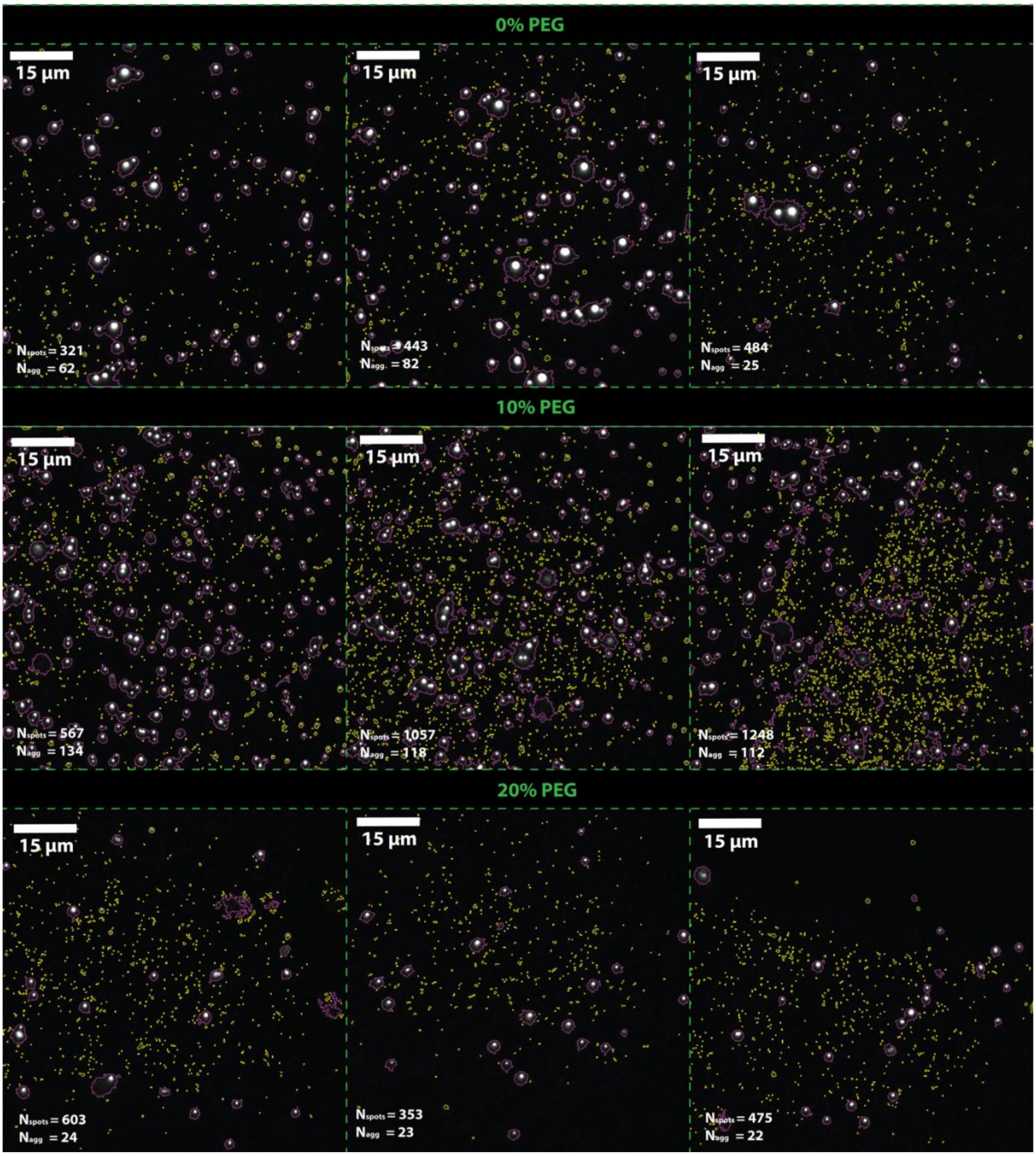


**Supplementary figure 6. Representative TIRF microscopy images of PE-Rhodamine labelled DPhPC liposomes.** Three representative images were chosen for each PEG3000 formulation (0% top, 10% middle, 20% bottom). Liposome spot ROIs were highlighted in yellow, and aggregate ROIs were highlighted in pink. The number of liposome spot ROIs ($N_{spots}$) and aggregate ROIs ($N_{agg}$) are listed on the bottom left of each representative image.

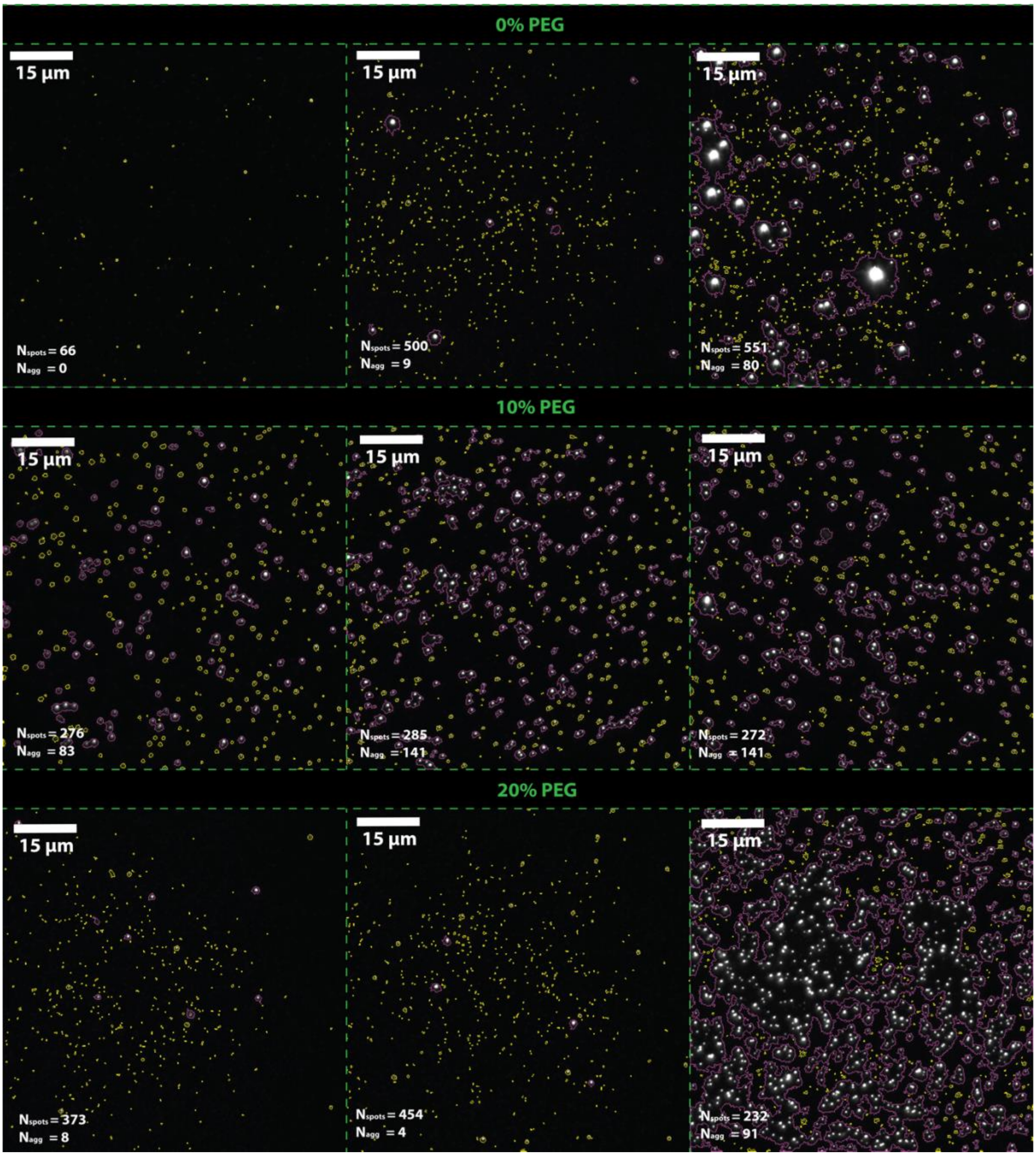


**Supplementary figure 7. Representative TIRF microscopy images of PE-Rhodamine labelled DOPE/DOPC liposomes.** Three representative images were chosen for each PEG3000 formulation (0% top, 10% middle, 20% bottom). Liposome spot ROIs were highlighted in yellow, and aggregate ROIs were highlighted in pink. The number of liposome spot ROIs ($N_{spots}$) and aggregate ROIs ($N_{agg}$) are listed on the bottom left of each representative image.

| DiD labelled liposomes | | | |
|---|---|---|---|
| **Lipid type** | **PEG3000-PE (%)** | $N_{spots}$ | $N_{agg}$ |
| DPhPC | 0 | 281 | 117 |
| | 10 | 142 | 166 |
| | 20 | 19 | 2 |
| DOPE/DOPC | 0 | 527 | 115 |
| | 10 | 5371 | 771 |
| | 20 | 1698 | 261 |
| **PE-Rhodamine labelled liposomes** | | | |
| **Lipid type** | **PEG3000-PE (%)** | $N_{spots}$ | $N_{agg}$ |
| DPhPC | 0 | 6810 | 533 |
| | 10 | 4469 | 604 |
| | 20 | 2724 | 101 |
| DOPE/DOPC | 0 | 6934 | 996 |
| | 10 | 3194 | 1186 |
| | 20 | 2548 | 628 |

**Supplementary table 4. Summary of total number of contour ROIs identified across all conditions.** For DiD labelled liposomes (top) and PE-Rhodamine labelled liposomes (bottom) the total number of contours was listed for liposome spot ROIs ($N_{spots}$) and aggregate ROIs ($N_{agg}$).
